# Modulation of decision-making latency by innate, learned and contextual factors in bumblebees

**DOI:** 10.64898/2026.02.08.704670

**Authors:** Romain Willemet

## Abstract

Foraging bee decision-making research has focused on choice determinants, and the variability and underlying causes of pre-choice latency remain understudied. Here, individual bumblebees (*Bombus impatiens*) were trained to associate one colored stimulus with a medium-value reward and another with a novel, higher-value reward. The experimental design consists of seven blocks, each containing four consecutive single-stimulus presentations followed by a forced binary choice. The latency to choose a stimulus and the type of choice during dual-choice trials were analysed. In dual-choice trials, bees in the yellow-high reward group showed a slower increase in high-reward selection than those in the blue-high group, suggesting persistent innate color bias. Response latencies for the low-reward stimulus systematically increased across blocks, indicating progressive devaluation. Early learning phases showed a temporary increase in response latency, extending previous findings on experience-dependent adjustments in acceptance thresholds. Latency in single-stimulus trials correlated with binary choice results, though choice proved a stronger indicator of preference than latency. Certain options elicited faster responses when presented with an alternative than when presented alone. Together, these findings support a deliberative model of bumblebee decision-making, in which pre-choice latency is modulated by innate preferences, associative learning, and immediate context.

## Introduction

An experimental subject’s decision to select a given option among alternatives involves a set of cognitive mechanisms that enable them to detect, identify, evaluate, and respond to that option (Green et al., 1966; Hammerstein & Stevens, 2012). Because these processes take time, it is possible to generate hypotheses about their nature by measuring and manipulating latency before choice. Such chronometric measurements have led to the development of a wide range of decision making models that have been used to evaluate hypotheses about decision making mechanisms and their neural correlates (e.g. Ratcliff & McKoon, 2008; Simen, 2012).

An important aspect of this research is to understand the extent to which decision making involves a direct comparison between options at the time of choice that goes beyond the automatic accumulation of evidence for each option (Bago & De Neys, 2017; Glöckner & Witteman, 2010; Salinas et al., 2014). Research on the chronometric properties of decision-making in honeybees and bumblebees; which are model species in animal cognition research (Bitterman, 1996; Chittka, 2017; Giurfa, 2025; Menzel, 2021; Muth et al., 2025), has led to recent developments on this issue. For example, MaBouDi et al., (2023) reinterpreted the “speed-accuracy trade-off” described in previous studies on bumblebees (e.g. Chittka et al., 2003) by demonstrating that the evidence threshold for accepting or rejecting a stimulus is not fixed. Bees tend to accept stimuli rapidly only when evidence accumulates quickly and strongly and, as a result, correct choices are, on average, actually faster than incorrect choices (MaBouDi et al., 2023). Thus, the relationship between speed and accuracy does not simply depends on the amount of evidence collected, as assumed by a simple deliberative account of speed/accuracy trade off, but also by the dynamics of the acceptation threshold. Another line of evidence seemingly against a deliberative model of decision making is the finding that honeybees evaluate options based on their reinforcement history with each option, rather than by comparing or ranking them during simultaneous choice (MaBouDi et al., 2020). In the study by MabouDi et al., however, the bees were trained with five options that differed in their probability of offering a reward. Such relatively high number of options may have added complexity when evaluating their respective properties (see also Benard & Giurfa, 2004).

Here, to further examine the extent to which choice involves deliberation, common eastern bumble bees (*Bombus impatiens*) were given a variant of the two-alternative forced choice task, one of the key task in decision making research, including in social bees (e.g. Giurfa, 2004; Shafir, 1994). The overarching goal was to better understand how decision-making latency evolves as bees learn the value of differently rewarding options when these options are presented alone or together. The literature on the Sequential Choice Model (SCM, Monteiro et al., 2020; Shapiro et al., 2008; Vasconcelos et al., 2010) provided useful context for this experiment. The SCM is a particular type of a race model of decision making, in which each option is processed independently until a decision criterion is reached in a parallel, race-like process (Heathcote & Matzke, 2022; Logan & Cowan, 1984). This model assumes no comparison between options at the time of choice. Instead, the SCM postulates that choice between known options presented simultaneously is the result of learned preferences during previous individual encounters with the various options (Shapiro et al., 2008). A remarkable prediction of this model is that the least preferred option should be generally chosen faster during dual choice trials compared to when it is presented alone. This arises because sampling latencies follow a distribution centered on a mean value; thus, only the shortest latencies associated with the least-preferred stimulus—which exhibits a longer mean duration—are likely to manifest behaviourally (Monteiro et al., 2020; Shapiro et al., 2008). Evidence supporting this prediction is mixed (Kacelnik et al., 2023). While no dedicated studies found a lengthening of latency during decision making, some reported no evidence for a shortening of latency for less-preferred options (e.g. Aw et al., 2012; Vasconcelos et al., 2013).

Examining the “deliberation” part of decision-making requires a measure of latency that reflects, as closely as possible, the cognitive process happening between the time a subject identifies a potential target and the start of the behavioural response to that target. To this end, large coloured stimuli were used here to facilitate visual search (Spaethe et al., 2001). Also, free flying toward a target, as is typical of bee visual cognition studies (e.g. Avarguès-Weber et al., 2010), involves a landing period that, while highly stereotyped (Baird et al., 2013; Reber et al., 2016; Srinivasan et al., 2001), can add noise to the latencies before landing. To address this issue, latency was recorded here as the time between a bee entered the arena (and almost simultaneously detected the target) and flew through a coloured target indicating a potential reward.

During a series of seven consecutive blocks, twenty bumblebees were individually presented with two types of stimuli, each associated with a particular reward and indicated by a specific colour. A block consisted of four trials where the stimuli were presented individually (twice each), and a dual choice trial in which they were presented simultaneously. The latency to choose a stimulus and, during dual-choice trials, the type of choice (high or low reward option), were recorded.

This experimental paradigm facilitates further investigation into the influence of innate colour preference on decision-making. Pilot observations indicated a pronounced bias toward blue stimuli. Previous research has demonstrated that innate colour preferences in bees significantly affect choice beyond initial exposures (Maharaj et al., 2019). However, the impact of such preferences on choice latency following repeated presentations—whether in isolation or simultaneous display—remains unclear.

Additionally, this paradigm enables examination of how reward quality influences choice. Bees typically exhibit a preference for higher-concentration sugar solutions within the 0-50% (w/w) range (Bailes et al., 2018; Konzmann & Lunau, 2014), yet the relationship between reward value and latency during foraging decisions is poorly understood. Higher-value options tend to elicit faster responses in many of the species tested so far, including humans (Shadmehr et al., 2019). The Sequential Choice Model (SCM) builds on this observation, positing that latency before choice in single-option trials serves as a robust indicator of preference (Kacelnik et al., 2023). Traditional research on bee cognition has primarily focused on choice outcomes, and empirical support for this effect remains limited. Evidence suggests bumblebees modulate flight speed based on reward value (Willemet, 2024), but the effect of multiple rewards and the learning of their properties on this relationship has not been established.

The present design also allows for testing a unique prediction of the SCM: that latency before choosing the least preferred option is reduced in dual-choice compared to single-choice trials. In fact, comparing latencies between single and dual-choice trials makes it possible to assess whether the presence of an alternative option influences choice latency, which would support a deliberative framework for decision-making.

## Methods

### Subjects

Two bumblebee colonies (*Bombus impatiens,* Biobest North America, Leamington, ON, Canada) were used in this experiment. Bees were provided daily with commercially available honeybee pollen deposited inside the nesting area. The colonies were housed in a experimental room surrended by windows providing enough natural light for orientation and flight. Two 5000K, 1600 lumens, Philips A19 led light bulbs were added to regulate luminosity throughout the day and facilitate video-tracking analysis.

### Apparatus

The flight arena was made of white plastic mesh and cardboard supported by a wooden frame, and covered by a black plastic mesh ceiling. A separation wall with a rectangular opening (height: 13 cm, width: 16 cm) at its centre could be installed to create two chambers (Figure 1.a).

**Figure 1.**
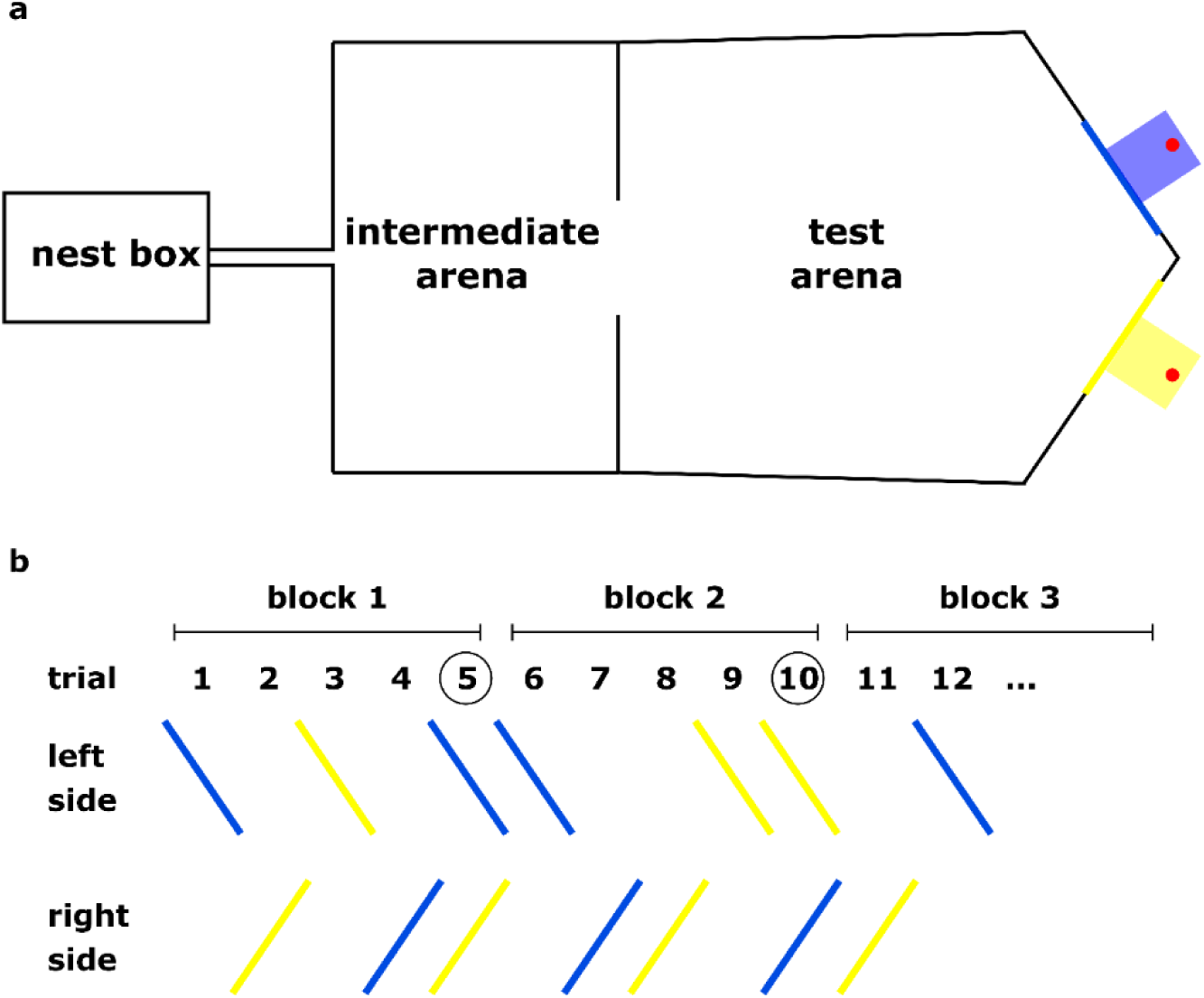
a. Top schematic view of the flight arena in a dual choice configuration. The intermediate arena is cuboid in shape (length: 60 cm, height: 56 cm, width: 40 cm). The back wall of the test arena is a 56 cm high wall with two 37 cm long sides at a 110° angle between each other. The test arena is thus 78 cm at its longest, and 56 cm at its shortest, from the separation wall, and its width is between 63 cm at the stimuli side and 60 cm at the separation wall. b. Example of a typical pattern of stimulus presentation. Trial numbers corresponding to dual trials are encircled.

On the stimulus side of the test arena, two openings on each side (centre at 12.5 cm from the middle of the wall and 27 cm high, diameter 8 cm) allowed the introduction of glass beakers. The sugar solution was deposited as large drops (allowing the bees to drink *ad libitum*) on small (3 cm diameter) red plastic disks that were changed between trials. To prevent odour cues from scent marks (Giurfa & Núñez, 1992) from influencing the results, glass beakers were cleaned with 91% isopropyl alcohol, rinsed with purified water, and dried after each trial. The arena and the nest box were connected via a GAWA type individual bee selector (Willemet, 2025) to reduce both the duration between bouts and the foraging experience variability between individual bees and bouts. The experiment was recorded with a GoPro Hero3 Black Edition camera (GoPro, Inc.); 720*1280; 120 fps), positioned 60 cm above the top of the arena.

### Stimuli

The target stimuli were yellow (RGB 255,255,0) and blue (RGB: 0,75,225) coloured rings 5.5 cm wide and 19 cm in diameter, which could be placed around the 8 cm opening of the glass beakers containing the sugar solutions. Stimuli of this size can be detected and discriminated in as little as 25 milliseconds in bumblebees (Nityananda et al., 2014), thereby minimising sensory processing time.

### Rewards

During pre-training and as the low-value reward, a 30% w/w sugar solution was used. The high-value reward consisted of a 45% w/w sugar solution. These concentrations were chosen during pilot experiments so that the bees would exhibit a strong preference for the high-value reward, while continuing to forage on the low-value reward.

### Procedure

#### Pre-training

During pre-training, the rewarding beaker was signaled by a grey ring, and its side randomely changed. Once several bees had started to forage autonomously, the separation wall was progressively raised. After this stage, the trained bees were individually marked using dots of acrylic paint, and selected for the experiment.

#### Experiment

The experimental design comprises seven blocks, each consisting of five trials (Figure 1.b). The first four trials of each block are single choice trials in which only one coloured stimulus and its associated reward is present in the arena. Each stimulus appears on the left and right side once per block. The order of colour and side of presentation are random. The fifth trial consists of a dual-choice trial in which both stimuli are presented simultaneously. The side of the low- and high-quality rewards during dual-choice trials is random between blocks.

Individual foraging bees performed these seven blocks in a single, continuous session. During each trial, bees could feed *ad libitum* on the sugar solution before going back to the nest. The bees were alternatively assigned to the blue-high/yellow-low (9 bees) or the blue-low/yellow-high (11 bees) reward configuration.

### Response variables

#### Choice

In dual-choice trials, the initial stimulus selected by bees was recorded, corresponding to either the low- or high-reward option.

#### Latency

Latency was recorded by watching the video recordings at low speed and coding the time between when the bee entered the test arena to when it entered the glass beaker.

Technical issues led to 8 latencies (out of 700) not being recorded. Latencies above 20 seconds, which involves other exploratory behaviours rather than choice, were excluded from the analyses (66 out of 692). Thirty-nine of these sixty-six instances happened during the first trial, in which bees often had to be forced to enter the stimuli after a 60 second threshold (by capturing them with a glass vial and introducing them inside the glass beaker containing the rewarding solution). In order to have one latency value per condition, the values from the two similar trials (high or low) within blocks were averaged.

### Statistical analysis

To analyse choice during dual choice trials and latency before landing, generalized linear mixed-effects models (package nlme, Pinheiro et al., 2019) were computed with the following fixed factors and their interaction: blocks, colour of high reward stimulus, type of trial (high, low, dual) and subject (bee) as random factor. Logistic regression were used to analyse prediction of choice based on latency in single choice trials. Sign tests were used to test for potential differences in latency differences between dual and single-choice trials. Block 1 was excluded from several analyses due to neophobia (see above). All analyses were ran using R Statistical Software (v4.2.1; R Core Team 2022).

## Results

Representing the evolution of latency over blocks reveals a combination of effects of colour of the stimulus associated with the high reward, blocks, and trial type (Figure 2).

**Figure 2.**
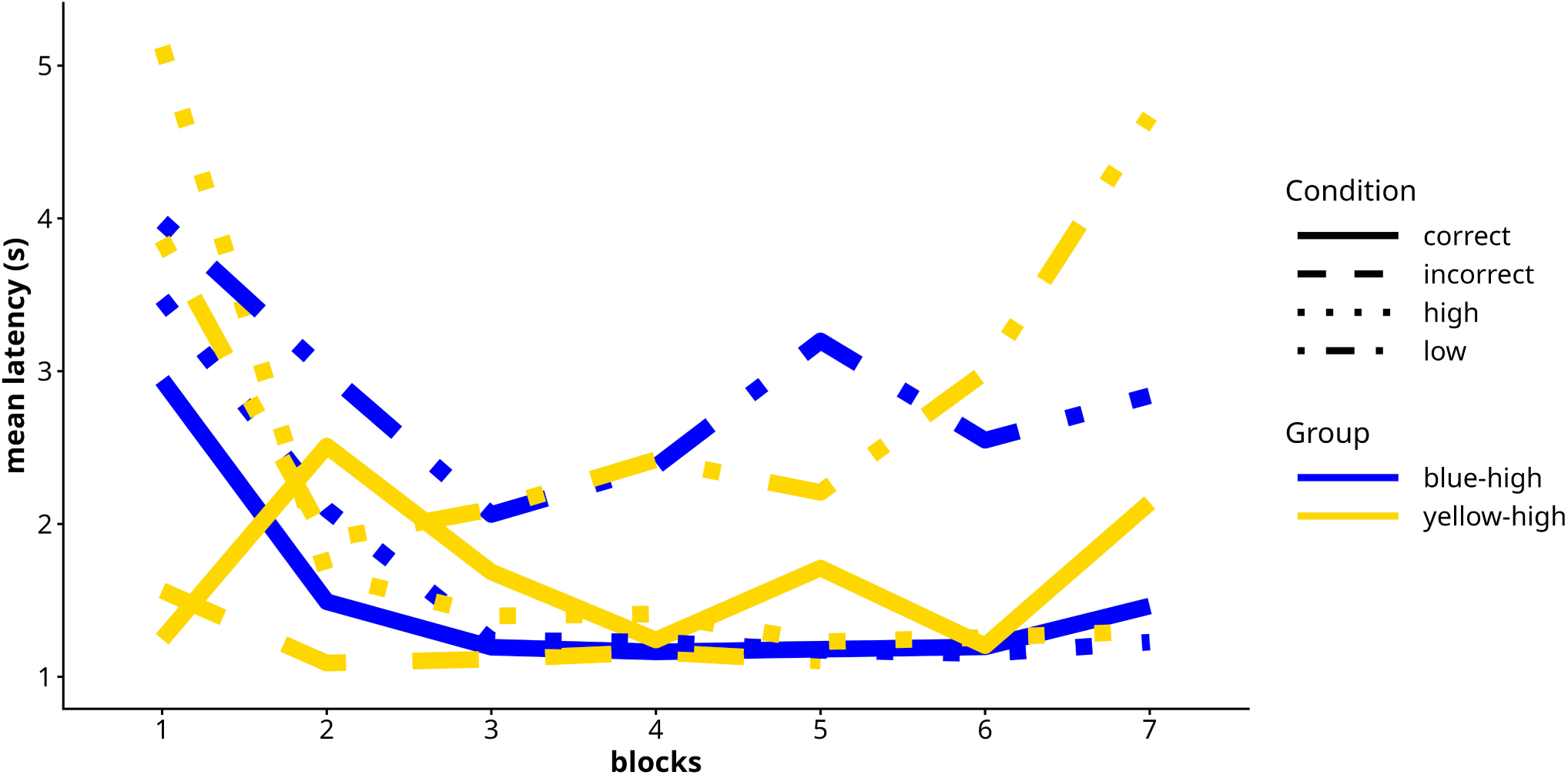
Average latency over blocks for blue-high and yellow-high groups during single choice trials (trial types high and low) and dual choice trials (“correct” when the bee choose the highest option, and “incorrect” otherwise). Only groups for which at least three datapoints were available are represented.

### 1. Influence of innate colour preference

During the first dual-choice trial, 7 out of 9 bees of the blue-high group, and 7 out of 11 bees of the yellow-high group choose the blue stimulus, a slight but not statistically significant preference for blue (binomial test: p-value = 0.1153). However, a lasting difference between groups exists in the proportion of bees chosing the highest reward during dual-choice trials (Figure 3). A GLMER analysis indicates a significant effect of colour (χ²(1) = 10.52, *p* = 0.001) as well as blocks (χ²(1) = 10.74, *p* = 0.001) on choice with blue-high bees rapidly reaching full blue choice, while yellow-high bees took longer to avoid blue despite its lower reward value.

**Figure 3.**
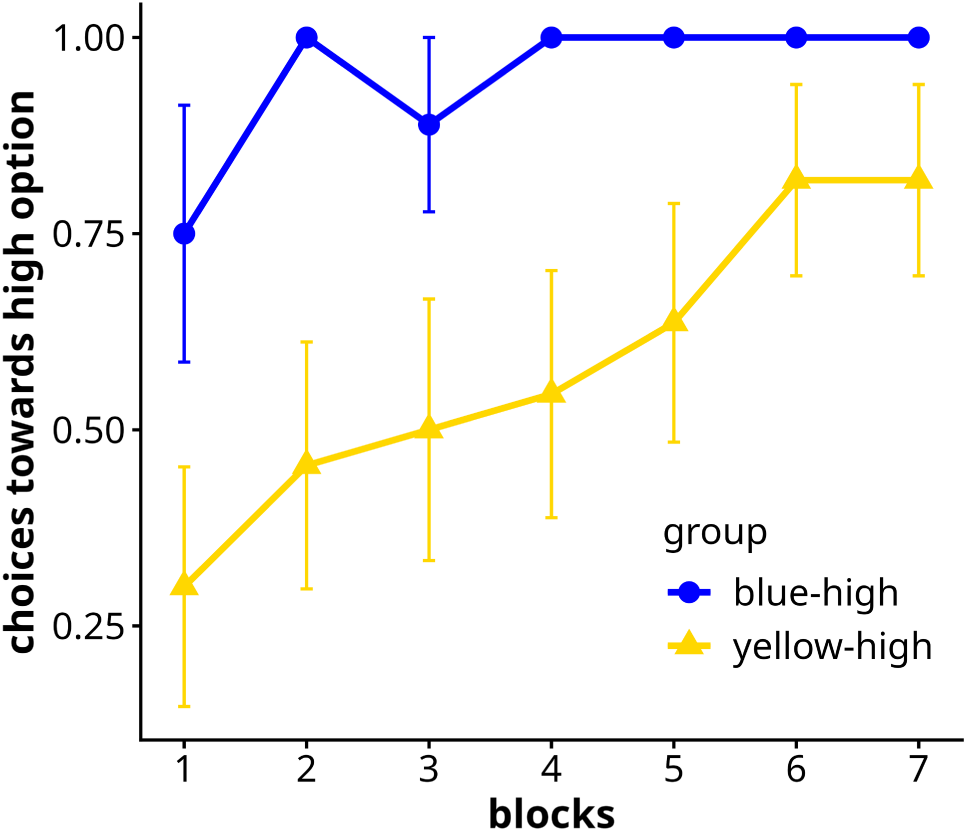
Proportion of choice (mean ± SE) during dual-choice trials towards the high option over blocks in blue- and yellow-high groups.

### 2. Influence of reward quality on latency in single choice trials

Linear mixed-effects models reveal a significant effect of condition (high or low reward) and interaction between condition and block on latency for bees in single choice trials for both the blue-high and yellow-high groups (p-values < 0.05) when block 1 is excluded. In both groups, the latency for low rewards increased over blocks, and the latency for high reward decreased and remained low after block 2 (Figure 2).

### 3. Influence of learning on latency

Bees from the blue-high group choosing blue tend to take longer before landing on the chosen stimulus than other groups in the first dual-choice trial (Figure 4). A Kruskal-Wallis rank sum test revealed a significant difference in latency across the groups when the two bees (a number too small for the test to be computationally valid) from the blue-high group choosing yellow are removed (χ2=6.4254, df=2, p=0.04). Pairwise Dunn tests indicate no significant differences between the groups of yellow-high bees choosing blue or yellow, but a significant difference between these two groups and the group of blue-high bees choosing blue (p-values < 0.05).

**Figure 4.**
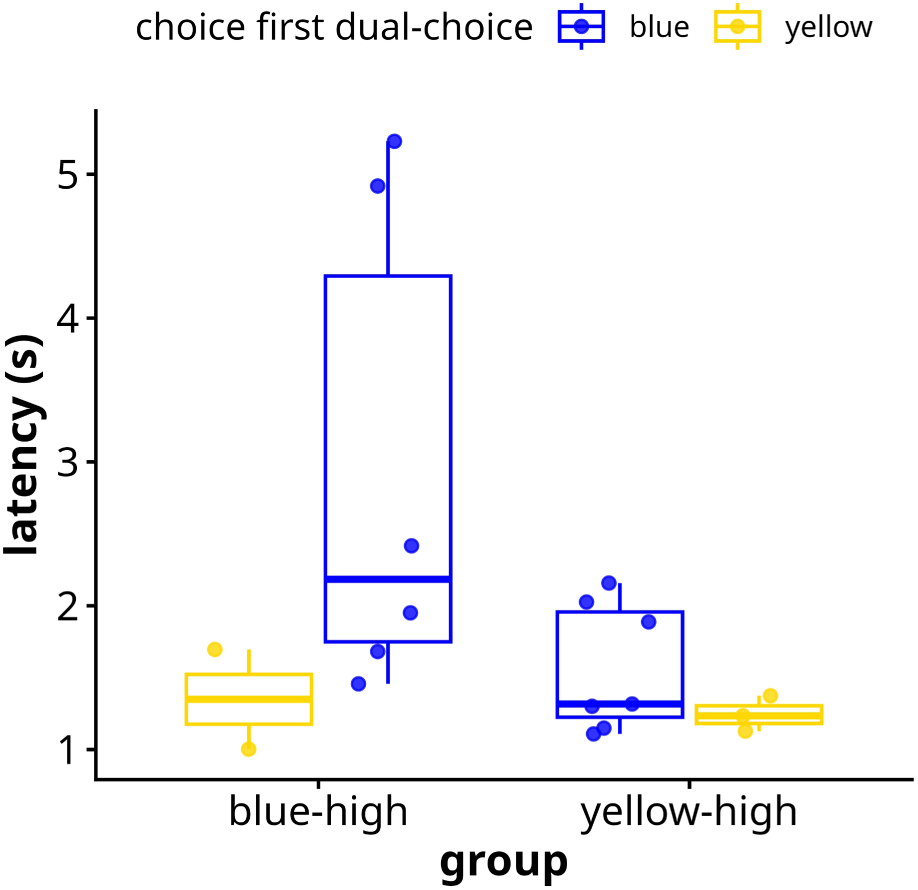
Latency (in seconds) before choice during the first dual-choice trial (block 1).

### 4. Comparing single and dual-choice trials

#### a. Does single-trial latencies predict decision-making during dual-choice trials?

The choice (high or low reward) during dual-choice trials can be predicted by the latency during the preceding single-choice trials 85% of the time (46 out of 54) in bees from the blue-high group (block 1 excluded). All but one of the failed predictions are from faster average latencies on yellow compared to blue during single trials. In the yellow-high group, predicting results during dual-choice trials based on the shortest average latency between the options during the four preceding single-choice trials yields a lower, albeit significative 65% of accuracy (logistic regression: 1.278, SE = 0.539, z = 2.372, p = 0.018). The prediction rate for the yellow-high group does not increase with blocks (glmer: Type II Wald chi-square tests χ²(5) = 4.88, p = 0.430, Figure 5).

**Figure 5.**
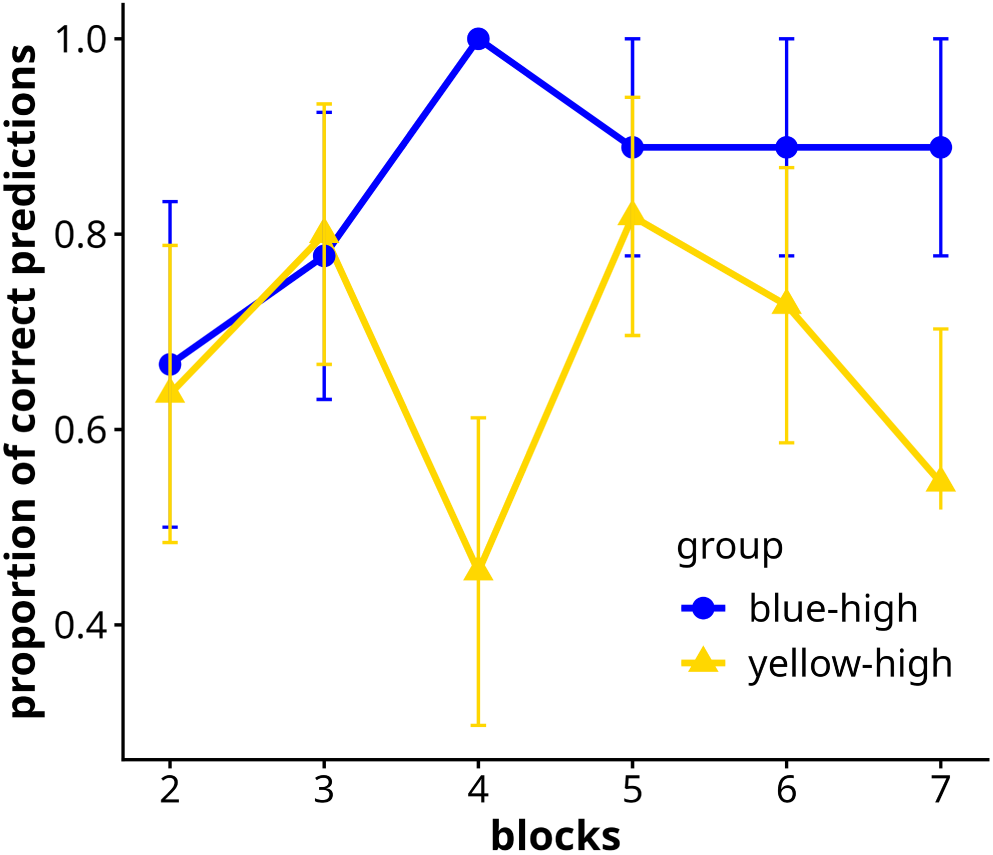
Proportion (mean ± SE) of dual-choice trials whose outcomes could be predicted by the option associated with the smallest latency during the preceding single-choice trials of that block.

#### b. Are latencies during dual and single-choice trials similar?

##### 1. Latency differences between trial types

To analyse whether the type of trial (single or dual-choice) influences latency before choice, the latency during choice was compared to the latency during the preceding single-choice trials for the option chosen. For example, if a bee from the yellow-high group chose blue during a dual-choice trial, the latency during choice in that trial is compared to the average latency during single-choice trials for blue. When data from blocks 2 to 7 are aggregated and the difference between the two latency values is examined (Figure 6), no difference between trial types is found for bees from the blue-high group (mean difference = -0.07 seconds, SD = 0.63, sign test (24|53): p-value = 0.583). Despite some values driving the average latency difference above zero, there is also no overall differences in latency between trial-types in bees from the yellow-high group chosing yellow (mean difference = 0.34 seconds, SD = 1.51, sign test (20|41): p-value=1). By contrast, latencies tend to be shorter during dual-choice trials compared to single choice trials for bees from the yellow-high group chosing blue during dual-choice trials (mean difference = -0.35 seconds, SD = 1.04, sign test (6|23): p-value=0.035).

**Figure 6.**
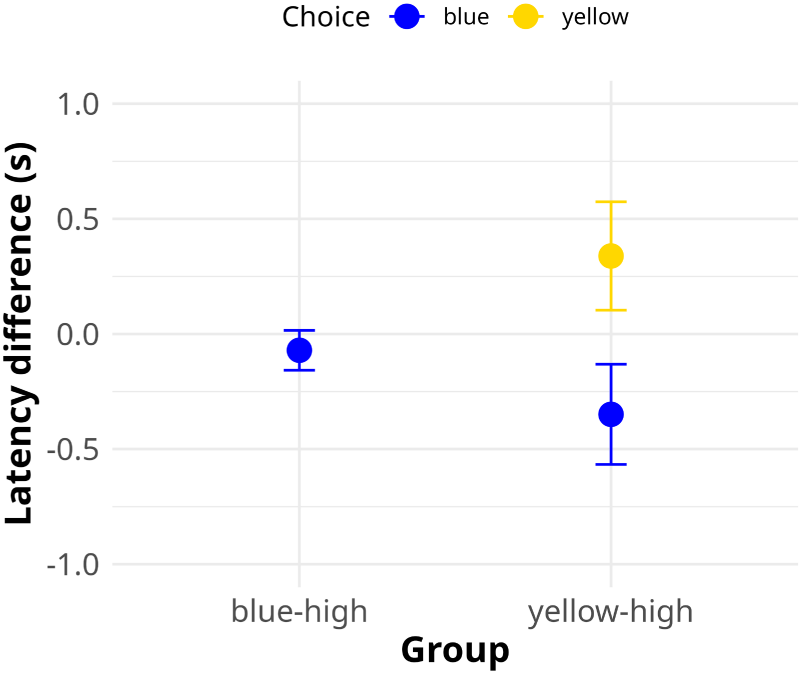
Latency differences (mean ± SE) between single and dual-choice trials for blue-high and yellow-high groups. The latency difference is calculated as the difference between the latency in dual-choice trials and the average latency in the two preceding single-choice trials corresponding to the colour chosen during dual-choice trial.

##### 2. Latency differences between preference groups and conditions

The analysis of latency differences between trial types shown above is limited by the lack of consideration of the individual preferences for each option. Since the SCM hypothesis of shortening of latency concerns the least preferred option, it is necessary to distinguish between groups of bees based on their level of preference for each option. That is, while the analysis above compared, for each group (blue-high and yellow-high), the latency during dual-type trials to the latency during single-choice trials of the color corresponding to the option chosen, the present analysis also considers groups based on their prefered option before choice. Their prefered option is determined by the shortest average latency during the two single-choice trials preceding the dual-choice trial. As the direction of the change, rather than the magnitude of the change itself, is the variable of interest here, and considering the effect of large latencies values highlighted by the previous analysis, a latency index was devised to represent the data (Figure 7). This latency index equals 1 when the latency during dual-choice trial is longer than the latency during single-choice trials, and –1 when it is shorter. Analysing the latency index using sign test suggests that no significative pattern or difference (shorter or longer) can be found in any of the groups, except for the group of yellow-high bees going faster on the blue stimulus during single-choice trials and choosing blue during dual-choice trials (sign test (3|15): p-value= 0.035).

**Figure 7.**
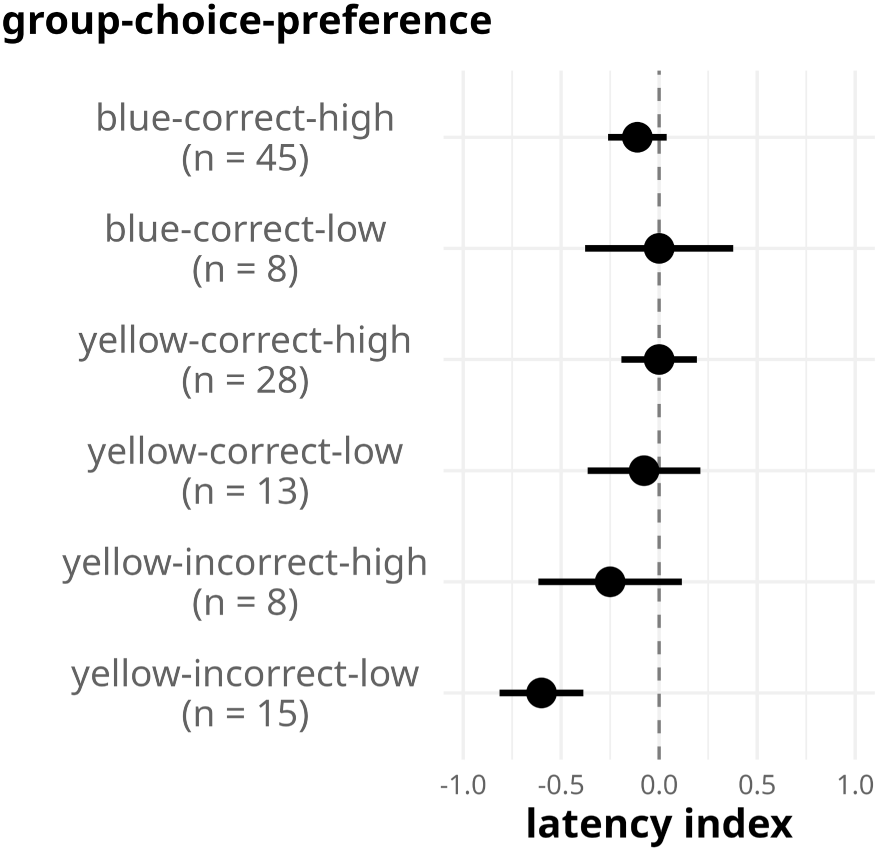
Latency index (mean ± SE) in groups of bees based on the initial group (blue or yellow-high), the response on the dual-choice trial (“correct” and “incorrect” have been used here instead of “high” and “low” rewards, to simplify the notation), and the fastest latency during the preceding single choice trials (low reward is yellow for the blue-high group, and blue for the yellow-high group). Negative latency index indicate shorter latencies during dual-choice trials compared to single-choice trials.

##### 3. Latency differences between groups during dual-choice trials

The analysis above suggests that groups based on the high-reward colour and their prefered colour differ in how they respond to a change in context (single vs dual-choice trials). The same groups can be compared in how fast they reach their target during dual-choice trials (Figure 8). Kruskal-Wallis test suggests significant differences between groups (chi-squared = 11.693, df = 5, p-value = 0.039). A follow-up Dunn test reveals a significative difference between the bees from the blue-high group that chose the blue (high) option during test depending on whether their fastest preceding single choice trials were yellow or blue (z-statistic = 1.766, p-value = 0.039). The group of bees from the yellow-high group that chose blue during dual-choice trials after having shorter latencies for blue during the preceding single-choice trials (denoted here as “yellow-incorrect-low”) were significantly faster than all other groups except the group of yellow-high bees choosing blue on dual-choice trials and going faster on blue stimuli compared to yellow during single-choice trials (p-values< 0.05).

**Figure 8.**
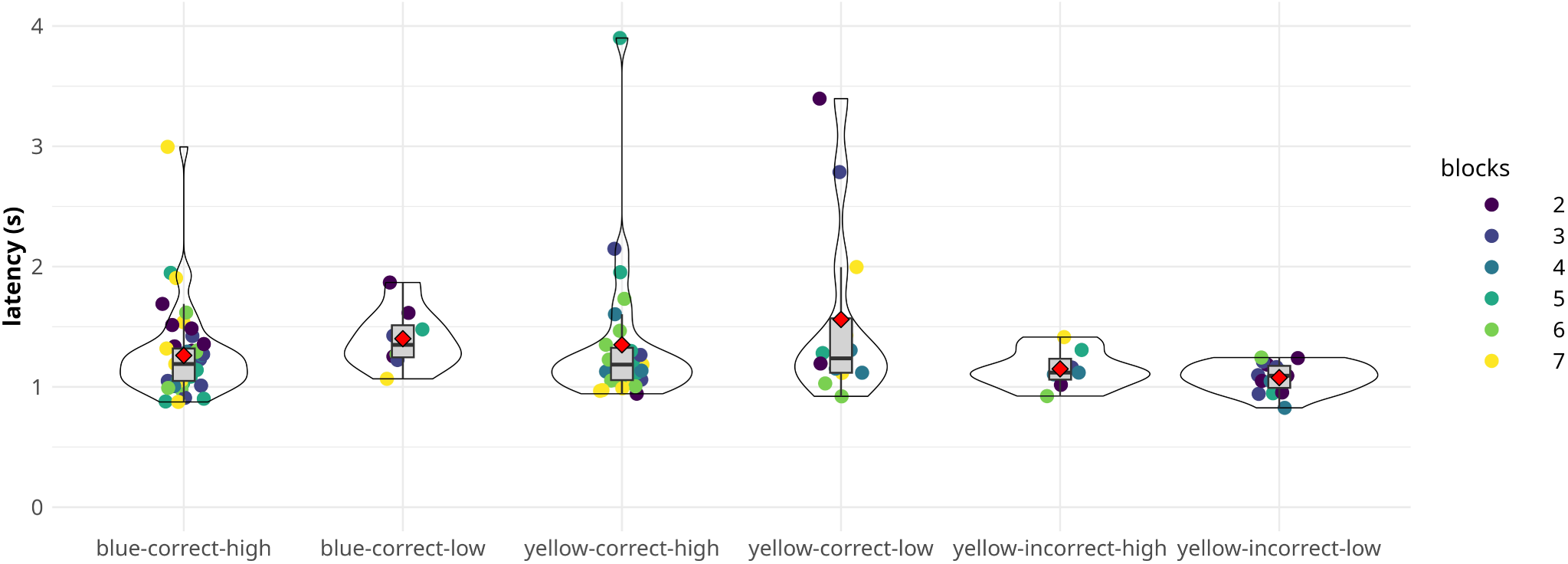
Latencies before choice in dual-choice trials for groups of bees based on the initial high-value reward (blue or yellow), the response on the dual-choice trial (“correct” and “incorrect” indicate “high” and “low” rewards), and the fastest latency during the preceding single choice trials. Red diamonds indicate the mean.

## Discussion

### 1. Innate preference has strong and lasting consequences

Innate colour preference, particularly towards the human blue range of colours, is a common factor in bee cognition research (Chittka & Wells, 2004). Differences have been reported between species (Balamurali et al., 2018; Heuel et al., 2024), and even between populations within species (Ings et al., 2009; Raine et al., 2006). Contrasting results exists for common eastern bumblebees regarding a preferential bias for blue (Forrest & Thomson, 2009; Muth et al., 2018), which may be partly due to the visual environment in which the stimuli are presented (Forrest & Thomson, 2009; Rivest et al., 2017). The present paradigm does not allow to measure innate preference directly, as bees already had two encounters with each stimulus before encountering them simultaneously. Bees were slightly, albeit not significantly so, more likely to attend to the blue stimulus during the first dual-choice trial (14 out of 20). Most remarkably, the potent preference for blue is visible in the distinct patterns of evolution of latency (Figure 2) and choice between options (Figure 3).

The pronounced effect observed aligns with previous findings indicating that innate preferences influence not only the initial tendency to sample a stimulus but also subsequent responses to changes in predictability and reward value (Liga et al., 2024; Maharaj et al., 2019; Waldron et al., 2005). Additionally, preferred colors have been shown to be more readily learned (Gumbert, 2000; Menzel, 1967), suggesting that innate preferences systematically shape the processing of stimulus-related information during cognitive tasks beyond the initial exposure. As the establishement of a preference towards an option is facilitated by the presence of aversive reinforcement (Avarguès-Weber et al., 2010), the fact that here both solutions were rewarded may help explain the long lasting effect of the blue preference.

### 2. Bees appear to slow down to evaluate options during learning

The extended latency observed in blue-high bees landing on blue during the first dual-choice trial (Figure 4), and the relatively long mean latencies for yellow-high bees landing on yellow in subsequent dual-choice trials (Figure 2) suggest that bees take longer to attend to a stimulus while learning its rewarding properties. These results are partially in line with those from a recent experience in rats using a comparable design consisting of dual-choice trials interseded between single-choice trials (White et al., 2024). White et al., (2024) reported longer decision-making latencies during the early phase of learning when rats encountered two differently rewarded stimuli simultaneously after initial individual exposure. Unlike the current results, the extended latencies in rats persisted through training, although they gradually diminished with learning (White et al., 2024). The discrepancy between studies likely stems from methodological differences rather than divergent cognitive mechanisms. For instance, rats underwent hundreds of trials over multiple days, whereas the present protocol consisted of 35 continuous trials given in a few hours.

The observed increase in decision-making latency during learning is consistent with established findings on information-seeking behaviour in bees. Bees modify their visual scanning patterns in response to the visual stimuli (i.e. active vision, MaBouDi et al., 2025), exhibit longer latencies when evidence acquisition is challenging (MaBouDi et al., 2023), and adapt their foraging strategies based on environmental difficulty (Dyer & Chittka, 2004; Ng et al., 2021; Perry & Barron, 2013; Spaethe et al., 2025). They also engage in “turn back and look” behaviour to reassess reward-associated locations (Lehrer, 1993, see also Guiraud et al., 2025).

The present results reinforce these observations, indicating that environmental unpredictability during learning directly affects pre-choice latency. This delay may arise from uncertainty about a specific option due to discrepancies between memory and stimulus predictions. Alternatively, it may reflect broader uncertainty regarding the relative values of available options. As bees begin to recognize differences between options, prolonged latency may serve to gather sensory information and identify the optimal choice. Another possibility is that increased latency stems from generalized uncertainty, where bees have not yet determined that options differ (e.g., that color signals reward value). Repeated mismatches between expectation and outcome—such as encountering a low-quality reward instead of a high-quality one—may heighten information-processing behaviour. Regardless of the mechanism, the data suggest that the acceptance threshold for a given option is elevated during learning, resulting in extended decision times.

### 3. Latency before choice as a measure of preference

The Sequential Choice Model posits that latency in single-choice trials reflects preference for an option (Kacelnik et al., 2023; Macías et al., 2021, see also Herrnstein, 1970). This study supports this assumption in the blue-high group, where dual-choice outcomes were largely predictable from single-choice latencies (around 85% of the times). However, there is more variability in the sampling latencies during single-choice trials than there is in choice during dual-choice trials (where blue was chosen nearly 100% of the time, Figure 3). This suggests that sampling latency may not be as strong an indicator of preference between options as choice between options presented simultaneously.

In the yellow-high group, single-choice latencies predicted dual-choice behaviour only 64% of the time. While this could be interpreted as reflecting the establishement of preference, the lack of linear evolution of predictability with blocks (Figure 5) suggests instead that several factors are competing against each others, even during later blocks. In fact, the discrepancy between the linear pattern of evolution of choice with blocks in yellow-high bees (Figure 3) and the lack of such a pattern in the predictability of choice based on single-choice trials (Figure 5) underscores that single-option latency is a more variable measure of valuation than choice in multi-option situations. The possible group differences in yellow-high bees selecting blue, depending on which option elicited faster responses in prior single trials (Figure 7), further suggest that single-option latency and multi-option choice offer complementary—not equivalent—measures of preference.

### 4. Effect of the presence of other options on latency

Bees in the blue-high group exhibited comparable latencies when entering the blue stimulus during both dual-choice and single-choice trials (Figure 6). Similarly, bees in the yellow-high group selecting yellow showed consistent latencies across trial types, though with greater variability. In contrast, yellow-high bees selecting blue during dual-choice trials responded faster when blue was presented alone (Figure 6). While the faster blue selection in yellow-high bees during dual-choice trials might appear to support the Sequential Choice Model (Kacelnik et al., 2023), bees exhibited shorter latencies for blue in single-choice trials compared to yellow in over 60% of these instances (15/24). This suggests that such “errors“—selecting the less rewarding option—may not be errors from the bees’ perspective, as they may inherently value blue over yellow.

Examining this in more detail by including the interaction between preference, trial type, and choice (Figure 7) reveals that yellow-high bees showing a preference for blue (based on preceding single-choice latencies) landed faster on blue when both options were presented. This effect was found in both relative and absolute terms, with this group demonstrating shorter latencies compared to most others (Figure 8). These findings contradict the SCM’s argument that the shortening of latency sometimes seen in multi-choice settings results from statistical sampling of the least preferred option’s latency distribution.

Instead, the results suggest a cumulative effect of attraction and rejection properties for each option. Yellow-high bees selecting blue in dual-choice trials may respond faster due to an innate bias towards blue, an aversion to yellow, and low perceived risk, given yellow’s high reward value. Alternatively, increased motivation when presented with two favorable options (blue due to innate attraction, yellow due to high reward) could explain the behaviour. The occurrence of rapid blue responses in early blocks supports the first hypothesis. However, bees can quickly associate a colored stimulus with high reward (Menzel, 1967), so the hypothesis of increased motivation in a low risk context is plausible. Either way, in this view, blue-high bees selecting blue responded quickly due to innate blue preference and yellow aversion, but not maximally fast initially due to higher perceived risk, as yellow was associated with lower reward. Finally, yellow-high bees selecting yellow may have showed slower latency reduction due to the conflict between reward value and initial color preferences.

The influence of additional options on bee decision-making may thus depend on the cost associated with each option. Armand et al. (2025) found limited evidence for the “decoy effect,” where unrewarded options increase preference for rewarded ones, including in earlier studies reporting the effect of additional options on choice (Latty & Trueblood, 2020; Shafir et al., 2002). In their experiment, small quantities (8 µL) of high-reward nectar (50%) per flower required bees to visit multiple inflorescences, reducing the cost of mistakes and potentially masking decoy effects. Here, bees responded faster when risk was low, indicating that lower mistake costs reduce behavioural inhibition. This aligns with findings that aversive stimuli enhance learning performance by reducing visits to unfavorable options (Avarguès-Weber et al., 2010).

### 5. Deliberation at the time of choice

The observation that bees in the yellow-high group exhibit faster responses to their preferred option when presented alongside a less preferred option contradicts a choice mechanism based solely on the expression of learned preferences. Instead, this result suggests a comparative evaluation of options at the time of choice, consistent with empirical evidence of deliberation in bee decision-making. For instance, Shafir & Yehonatan (2014) used a two-alternative forced-choice proboscis-extension conditioning paradigm to investigate how bees evaluate options when multiple reward dimensions (delay, duration, sucrose concentration) are manipulated. After training honeybees to associate two odors with rewards differing along one fixed dimension, both odors were presented simultaneously. Although a secondary reward dimension was manipulated similarly for both options, its variation did not affect choice, suggesting bees compared options based on their relative difference along the fixed dimension rather than absolute value.

The reduced latency observed here during dual-choice trials, which implies an interaction between options, occurs as preferences for the highest-value option are being established. The SCM, however, focuses on explaining choice mechanisms after preferences have already been learned (Shapiro et al., 2008). While several mechanisms of decision making may be involved depending on the level of experience (Kelly & Barron, 2022), choice situations in natural settings are likely to be constantly evolving situations. This is because choice is influenced by the subject’s state (Mayack & Naug, 2015; Pompilio & Kacelnik, 2005), environmental conditions, and alternative profitability (Hemingway et al., 2024; Shapiro et al., 2008). A compelling aspect of decision-making deliberation is the modulation of behavioral vigor based on expected reward (Shadmehr et al., 2019). Proponents of the SCM argued that such “delay to act” should be minimal when only one option is present due to the increased cost involved (Kacelnik et al., 2011). The speed of a behaviour is, however, rarely maximal, and animals, including humans adjust their behavioural vigor to the value of the options present in the environment (Sukumar et al., 2024). Here, sampling latency for the 30% w/w solution increased as bees gained experience with the more rewarding 50% w/w option, mirroring other findings in artificial foraging settings (Nityananda and Chittka, 2021). Although behavioural vigor in bees have seldomely been examined in the context of decision making, there is some evidence that bees also adjust their flight speed based on the expected reward (Willemet, 2024). The same adjustment mechanism between available options and environmental value may explain latency differences in single-choice situations.

The consideration of environmental context during choice raises questions about how multiple options are evaluated. The SCM assumes that deliberative choice involves simultaneous assessment of present options (Kacelnik et al., 2011). However, deliberation may also occur between the value of a given option and that expected from the environment (which can include other options). The vicarious trial and error literature in rats provides the clearest behavioural and neurological evidence for serial evaluation of option values (Redish, 2016). This process is rapid when the different values are known, but prolonged when elements are novel or variable. The sequential nature of decision-making and deliberation at choice are thus not mutually exclusive (Orquin & Mueller Loose, 2013). This process involves separating selective attention from behavioural choice, a phenomenon observed in humans (Moerel et al., 2024) and bees (Paulk et al., 2014).

A serial mechanism would align with the SCM argument that single-option decision mechanisms extend to multiple-choice scenarios (Kacelnik et al., 2011), albeit with deliberation. In that case, the primary difference between single- and multiple-choice environments is that available alternatives modify the environmental representation, for example by altering the cost of attending to, or searching for, a specific option. Interestingly, however, most insect studies favour parallel valuation models, with a winner-takes-all mechanism. Computational models using neuroanatomical data suggest inhibitory coupling between evidence accumulators is necessary to explain bee and fly behaviour (Barron et al., 2015; Groschner et al., 2018). These studies focused on olfactory processes, which may differ from visual ones. Inhibitory coupling appears essential to model acceptance and rejection behaviour in bees landing on a single visual reward (MaBouDi et al., 2023), though this model focuses on single-option evaluation (acceptance/rejection) of single flowers attented to sequentially.

More research using visual environments where options are available simultaneously is required to answer the degree of parallelism in bee decision-making. Social bees provide a particularly compelling model for investigating how alternative options alter the environmental representation during reward valuation, given interspecific variation in capacity for parallel visual attention. Compared to bumblebees, honeybees have reduced capacity for evaluating concurrently presented visual stimuli (Morawetz & Spaethe, 2012). This suggests that the availability of multiple options may differentially influence their decision-making. Supporting this, honeybees appear to employ distinct strategies compared to bumblebees in similar Matching-to-Sample tasks, seemingly being less influenced by the sample stimuli (Willemet, 2026). A comparison of both species would enable systematic investigation of the cognitive effects of restricted visual attention.

Free-flying bees are also particularly valuable for studying decision-making because pre-choice latency is embedded in foraging motor actions. Decision models often include a “non-decision time”, before the process is initiated (Logan et al., 2014). This has been criticized from a theoretical and empirical perspective (Palmer et al., 2025; Weindel et al., 2022). In bees, this phase is integrated with flight behaviour, providing opportunities to explore the subtleties of foraging decision-making and compare them with other model species (MaBouDi et al., 2023; Willemet, 2024).

More generally, while bee decision-making research has historically emphasized choice outcomes, the current findings contribute to a growing literature (e.g. Chittka & Spaethe, 2007; MaBouDi et al., 2023, 2025) demonstrating that pre-choice latency in foraging bees is a valuable research tool for studying the mechanisms behind decision-making.

## Acknowledgements

The author thanks Ellouise Leadbeater for helpful comments during manuscript development.

